# Interleukin-27 is antiviral at the maternal-fetal interface

**DOI:** 10.1101/2025.04.28.651113

**Authors:** Madeline S. Merlino, Briah Barksdale, Seble G. Negatu, Rebecca L. Clements, Taylor Miller-Ensminger, Alexandra H. Lopez, Sneha Mani, Monica N. Mainigi, Christopher A. Hunter, Kellie A. Jurado

## Abstract

Congenital viral infections can have severe consequences for pregnancy and fetal outcomes. Remarkably, the fetal-derived placenta serves as a robust barrier to infection through meticulous regulation by immune effectors and a diverse repertoire of cytokines. Yet, the regulatory roles of many cytokines remain undefined at the maternal-fetal interface. Interleukin 27 (IL-27) is a highly expressed cytokine in the placenta whose functional consequence during congenital infection is unknown. Here, we utilized trophoblast organoids (TO) derived from primary human placentas and a mouse model of congenital viral infection to uncover the functional role of IL-27 signaling during pregnancy. We show that TOs constitutively express IL-27 and its receptor, IL27RA, and demonstrate that IL-27 signaling restricts Zika virus (ZIKV) infection of TOs. Through bulk RNA-sequencing of TOs in the absence and presence of IL-27 signaling, we demonstrate IL-27-mediated upregulation of antiviral genes. Finally, we show that IL-27 signaling is critical within the context of congenital murine ZIKV infection, as IL-27 restricts placental ZIKV burdens and protects against pathologic fetal outcomes early in gestation. These findings collectively demonstrate a novel role for IL-27 in the placenta and establish IL-27 as an innate antiviral defense at the maternal-fetal interface during congenital viral infection.

## INTRODUCTION

During the critical developmental period of pregnancy, there are substantial consequences to infection. Despite complex immune regulation at the maternal-fetal interface, a subset of pathogens can evade host immune responses, cross the placental barrier, and establish congenital infection in the fetus.^1^ Zika virus (ZIKV), a mosquito-borne, positive-sense RNA flavivirus, is one example of a congenital viral pathogen.^2^ Depending on the time of infection, vertical transmission of ZIKV can cause adverse pregnancy outcomes such as microcephaly, preterm birth, and even fetal demise.^3–5^ As congenital ZIKV can have a detrimental impact on fetal health, there is a critical need to identify antiviral effectors at the maternal-fetal interface.

The fetal-derived placenta serves as an effective barrier to congenital viral infections through its constitutive expression of antimicrobial cytokines.^6^ For example, Type I interferons (IFNs) are a family of innate antiviral cytokines that are known to restrict viral replication through the rapid induction of IFN-stimulated genes (ISGs).^7^ However, while type I IFN (IFNα, ifNβ) signaling is essential for healthy gestation, the classical antiviral state driven by type I IFNs can also disrupt placental development and mediate fetal pathology during congenital ZIKV infection.^8,9^ Yet, the closely related type III IFNs (IFNλ) are constitutively produced by the placenta and have been shown to restrict ZIKV infection without causing fetal pathology.^10–12^ While the role of type III IFNs in placental immunity is well established, it is unknown whether other cytokines perform similar antiviral functions at the maternal-fetal interface.

Interleukin 27 (IL-27) is a heterodimeric cytokine that is produced by extravillous trophoblasts and syncytiotrophoblasts in the placenta throughout gestation; yet, the functional consequence of IL-27 signaling during pregnancy is unknown.^13^ In other contexts, IL-27 is a potent modulator of both innate and adaptive immune responses and has direct antiviral activities.^14,15^ Interestingly, a study by Kwock *et al.* found that IL-27 induces antiviral gene expression in a STAT1-dependent manner to confer protection against ZIKV infection in human keratinocytes.^16^ Together, these findings led us to ask whether IL-27 contributes to antiviral immunity at the maternal-fetal interface.

Using a human trophoblast organoid model, we reveal that IL-27 signaling is antiviral in fetal cells of the placenta. We then define the transcriptional profile of IL-27-stimulated trophoblast organoids through bulk RNA-sequencing and demonstrate IL-27-mediated antiviral gene expression in fetal trophoblasts, thus uncovering possible mechanisms of viral restriction by IL-27 at the maternal-fetal interface. Finally, we show that IL-27 signaling is critical within the context of congenital murine ZIKV infection, as IL-27 restricts placental ZIKV burdens and is protective against pathologic fetal outcomes. Together, these findings establish IL-27 as an innate antiviral defense at the maternal-fetal interface and highlight its potential for combating congenital ZIKV infections and supporting healthy fetal outcomes.

## RESULTS

### Primary human trophoblast organoids constitutively express IL-27 and IL-27RA

Prior studies indicate that IL-27 and its receptor, IL27RA, are expressed in the placenta throughout gestation.^13^ We generated primary trophoblast (TO) and decidual (DO) organoid lines from first trimester (6-12 post-conception weeks) human placental tissues according to previously established methods and initially sought to confirm IL-27 and IL27RA expression in these models (**Supplemental Table 1**).^17–19^ Prior to experimentation, TOs and DOs were grown in Matrigel to support self-organization of 3D structures and were routinely passaged every 6-8 days, with media changes every 2-3 days. We observed that our first trimester TO and DO cultures recapitulated the morphology of previously established TO and DO lines and exhibited distinct TO and DO structures that could be visualized via brightfield microscopy (**Figure 1A; Supplemental Figure 1A**).^17–19^ Additionally, purity of TO and DO cultures was validated prior to experimentation via over-the-counter pregnancy test strips, which detected a presence or absence of secreted trophoblast-specific hormone human chorionic gonadotropin (hCG) in TO- and DO-conditioned media, respectively (**Supplemental Figure 1B**).

**Figure 1:**
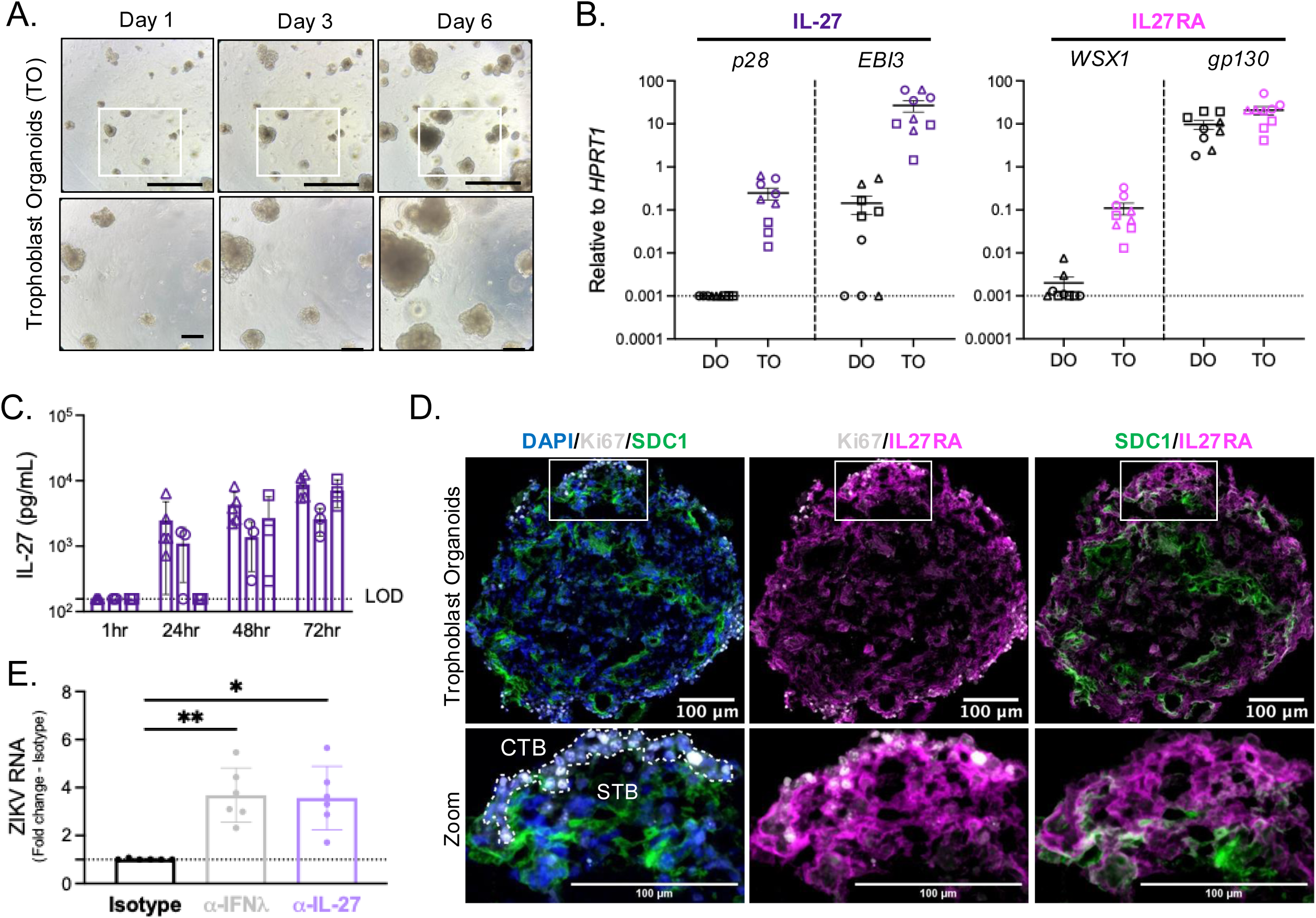
Primary human trophoblast organoids constitutively express IL-27 and IL-27RA. **A.** Representative brightfield images of single trophoblast organoid (TO) culture at 1-, 3-, and 6-days post-passage. 5X magnification, scale bar 1000μm (top). Insets denote area captured at 10X magnification, scale bar 200μm (bottom). **B.** Expression of IL-27 cytokine (*p28, EBI3*) and receptor (*WSX1, gp130*) subunit genes by decidual organoids (DOs) and TOs at day 6 post-passage as determined by quantitative RT-PCR, relative to housekeeping gene *HPRT1*. Graphs represent 3 experimental replicates from 3 biological donors (circles, squares, and triangles) for TOs and DOs. **C.** Conditioned media were collected from TO cultures at 1-, 24-, 48-, and 72-hours post-passage and secreted IL-27 was measured via human IL-27 ELISA. Graph represents 3-5 experimental replicates from 3 biological TO donors (circles, squares, and triangles). **D.** Representative immunofluorescent images of TOs at day 6 post-passage. 10μm TO cross-sections were stained for DAPI (blue), Ki67 for cytotrophoblasts (white), SDC1 for syncytiotrophoblasts (green), and IL27RA (magenta). White box indicates zoom inset and white dashed line demarcates regions of cytotrophoblasts (CTB) and syncytiotrophoblasts (STB). 20X magnification, Z-stack, merged. Scale bar, 100μm. Images representative of 3 biological TO donors. **E.** TOs were treated with either isotype control, α-IL-27 or α-IFNλ antibodies at days 3 and 5 post-passage. At day 5 post-passage, TOs then were infected with 10^5^ PFU Cambodian ZIKV (ZIKV-CAM) or PBS (Mock) for 24 hours. TO cultures were then collected at day 6 post-passage for RNA analysis. ZIKV viral RNA expression in TOs as determined by quantitative RT-PCR relative to housekeeping gene *HPRT1*, normalized to isotype control treated organoids. Graph represents 6 independent infections from 1 TO donor. Statistical analysis performed with Kruskal-Wallis ANOVA, *p<0.05, **p<0.01.

We first assessed the expression of IL-27 subunits (*p28, EBI3*) and IL27RA subunits (*WSX1, gp130*) in unstimulated TO and DO cultures via quantitative RT-PCR. We found that both TOs and DOs expressed high levels of transcripts for the shared cytokine and receptor subunits *EBI3* and *gp130;* however, mRNA expression of unique IL-27 and IL27RA subunits *p28* and *WSX1* was only observed in TOs (**Figure 1B**). To corroborate this finding, we measured secreted IL-27 protein in conditioned media from unstimulated TO cultures at various timepoints via enzyme linked immunosorbent assay (ELISA). We show that IL-27 is constitutively produced by TO cultures beginning as early as 24 hours post-passage, with increasing production as TOs grew in culture (**Figure 1C**).

To determine which cells within the multicellular TO are capable of responding to IL-27, we performed immunofluorescence microscopy for specific trophoblast markers and IL27RA. Ki67 staining identified highly proliferative cytotrophoblasts (CTB) at the periphery of TOs, while SDC1 staining identified multinuclear syncytiotrophoblasts (STB) at the center of TOs, thus demonstrating the inward-facing apical surface and “inside-out” orientation that is characteristic of TOs and other three-dimensional organoid models (**Figure 1D**).^18,20^ We observed strong IL27RA expression within CTB regions of TOs, with only some STB regions expressing IL27RA (**Figure 1D**; magenta). Together these findings indicate that IL-27 and IL27RA signaling machinery is intact in the TO model, thus validating TOs as an appropriate model to study IL-27 and IL27RA signaling in the human placenta.

We next sought to determine whether IL-27 signaling regulates congenital ZIKV infection in trophoblasts. Briefly, TO cultures were pretreated with isotype control or IL-27-neutralizing antibodies prior to infection with Cambodian ZIKV (**Supplemental Figure 2**). Additional TO cultures were pretreated with IFNλ-1 and IFNλ-3 neutralizing antibodies prior to ZIKV infection as a positive control, since IFNλ has been previously shown to regulate ZIKV infection of fetal trophoblasts through autocrine signaling.^10–12^ By measuring viral RNA via quantitative RT-PCR, we showed that TO cultures were susceptible to ZIKV infection, and that neutralization of IL-27 or IFNλ led to similar increases in TO viral RNA levels relative to control organoids (**Figure 1E**). These data indicate that IL-27 signaling restricts ZIKV infection of fetal trophoblasts similarly to IFNλ, and thus IL-27 is a critical mediator of antiviral immunity in the placenta.

### IL-27 induces antiviral gene expression in trophoblast organoids

Given the functional IL-27 signaling circuit in TOs and its known capacity to restrict viral infection in other contexts, we next sought to define IL-27-mediated gene expression changes in TOs. Since TO cultures are constitutively producing and responding to cytokines such as IL-27 and IFNλ (**Figure 1B-C**), we first aimed to quantify basal antiviral gene expression levels in TO cultures.^21^ To capture the basal TO state, TO cultures were treated with vehicle (PBS) and isotype control antibody (**Figure 2A; “Isotype”**). To broadly mitigate signaling downstream of cytokine receptors and capture an “unstimulated” TO state, additional TO cultures were treated with Janus-kinase inhibitor Ruxolitinib (Rux) and isotype control antibody (**Figure 2A; “Rux”**). We then measured the expression of common antiviral genes previously shown to be associated with IL-27 signaling (*MX1*, *OAS2* and *IFIT1*) in Isotype- and Rux-treated TOs via quantitative RT-PCR.^16,22^ As expected, we observed a strong basal expression of *MX1, OAS2*, and *IFIT1* in Isotype-treated TOs, with antiviral gene expression largely inhibited following Rux treatment (**Figure 2B**). Importantly, the antiviral gene expression observed in Isotype-treated TOs is reflective of all cytokines expressed by TOs in their basal state, including IL-27 and IFNλ. To specifically evaluate IL-27-mediated changes in antiviral gene expression independently of IFNλ, TO cultures were pretreated with IFNλ-1 and IFNλ-3 neutralizing antibodies followed by stimulation with recombinant human IL-27 (**Figure 2A; “IL-27”**). Conversely, to demonstrate IFNλ-mediated changes in antiviral gene expression independently of IL-27, additional TO cultures were pretreated with IL-27 neutralizing antibody followed by stimulation with recombinant human IFNλ1/3 (**Figure 2A; “IFNλ”**). Isotype- and Rux-treated TO cultures were also included as positive and negative controls, respectively (**Figure 2A**). We found that IL-27 stimulation of TOs in the absence of IFNλ significantly upregulated the expression of antiviral genes *MX1*, *OAS2* and *IFIT1*, relative to Rux-treated TOs (**Figure 2C**). Additionally, we observed no significant difference in the relative antiviral gene expression levels of IL-27-stimulated TOs and Isotype-treated TOs, which reflect gene expression changes downstream of all constitutively expressed cytokines. These results suggest that IL-27 has the capacity to induce antiviral gene expression in TOs independently of IFNλ, and that constitutive IL-27 signaling contributes to the robust basal antiviral state of TOs.

**Figure 2:**
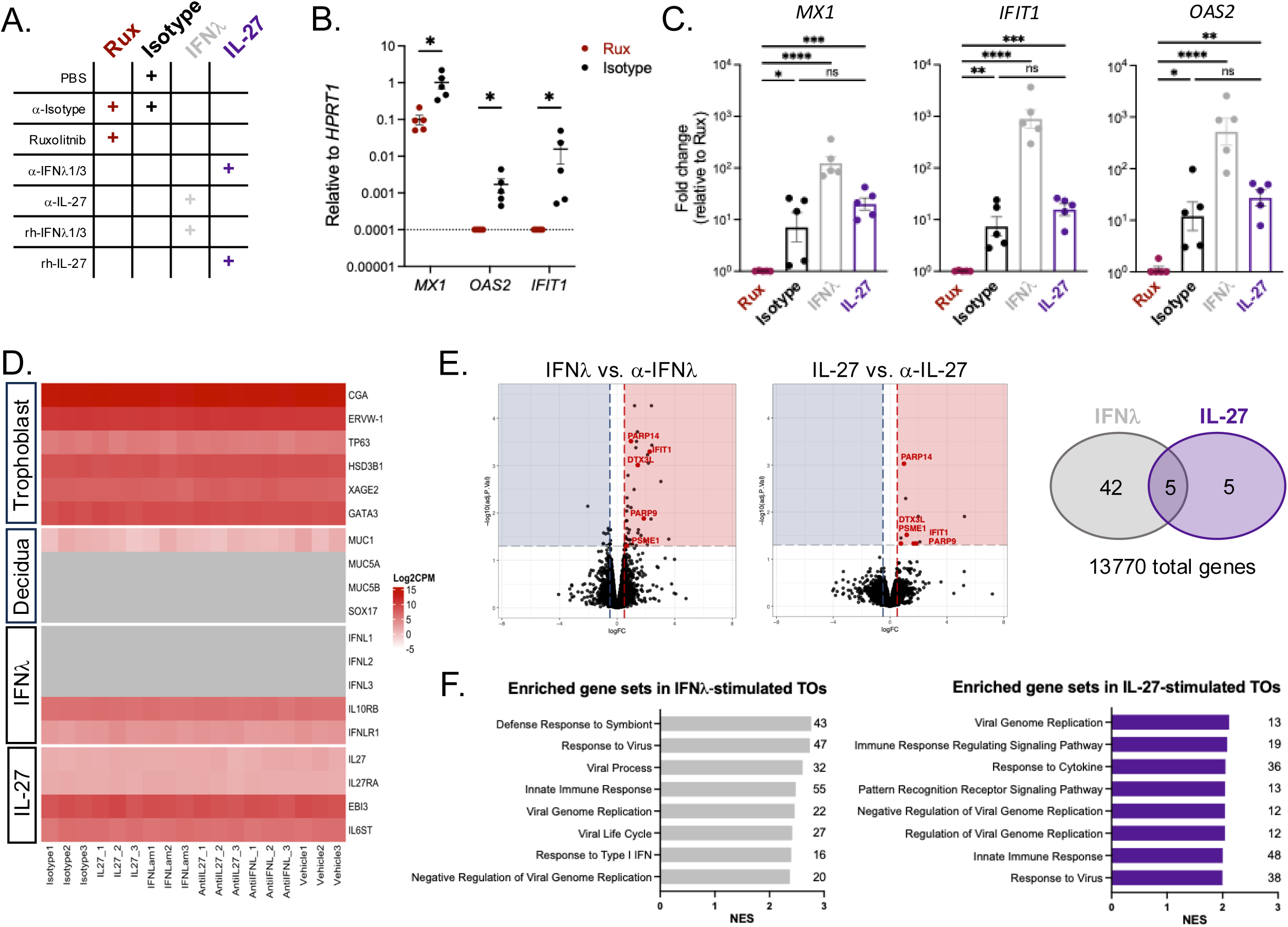
IL-27 induces antiviral gene expression in trophoblast organoids. **A.** Table of experimental conditions for antiviral gene expression analyses in **Figures 2B-C**. **B.** Expression of antiviral genes *MX1, OAS2,* and *IFIT1* by TOs as determined by quantitative RT-PCR, relative to housekeeping gene *HPRT1*. Graphs display 5 experimental replicates from 1 TO donor. Statistical analysis performed with multiple unpaired t-tests (Mann-Whitney), *p<0.05. **C.** Expression of antiviral genes *MX1, OAS2,* and *IFIT1* by TOs as determined by quantitative RT-PCR, relative to Rux. Graphs display 5 experimental replicates from 1 TO donor. Statistical analysis performed with Kruskal-Wallis ANOVA *p<0.05, **p<0.01, ***p<0.001, ****p<0.0001. **D.** Heat map depicting expression (log_2_(counts per million)) of common markers associated with the placenta, decidua, IL-27 and IFNλ signaling as determined by bulk RNA-sequencing of TOs. Dark red indicates high gene expression, light red indicated low expression, and gray indicates no detectable reads. **E.** Volcano plots depicting differentially expressed genes (DEGs) in IL-27-stimulated TOs compared to α-IL27 (10 upregulated DEGs) and IFNλ-stimulated TOs compared to α-IFNλ (47 upregulated DEGs). Vertical dashed lines on volcano plot represent a log fold-change cut-off of 0.5 and the horizontal dashed line represents an adjusted p-value cut-off of 0.05. The Venn diagram (right) depicts the total upregulated DEGs in the IL-27- and IFNλ-stimulated conditions, with the 5 shared genes highlighted in red text on the volcano plots. **F.** Select upregulated pathways from gene set enrichment analysis (GSEA) of bulk RNA sequencing data for IL-27- and IFNλ-stimulated TOs. Log fold-change values were calculated by comparing IL-27-stimulated vs. α-IL27 TOs and IFNλ-stimulated vs. α-IFNλ TOs. All GSEA analyses were then performed using a log fold-change cutoff value <0.5. Table lists upregulated pathways for IL-27-stimulated TOs by normalized enrichment score (NES). Figure displays top 8 enriched pathways by order of normalized enrichment score (NES). Numbers at end of bars indicate total number of genes associated witheach term.

We next aimed to further define the transcriptional impact of IL-27 signaling in TOs and identify possible mechanisms of viral control in an unbiased manner. TO cultures were pretreated with either IL-27-neutralizing, IFNλ-neutralizing, or isotype control antibodies for 72 hours prior to collection. Simultaneously, additional TO cultures were stimulated with either recombinant human IL-27, recombinant human IFNλ, or vehicle control (PBS) for 24 hours prior to collection. RNA was then isolated from all TOs and bulk RNA-sequencing was performed. We confirmed that TO cultures transcriptionally recapitulate the placental tissue of origin and express various placental markers (*ERVW-1, HSD3B1, XAGE2, GATA3*), including those specifically associated with STBs (*CGA*) and CTBs (*TP63*) (**Figure 2D**).^18^ As expected, we observed little-to-no expression of decidual markers *MUC1, MUC5A/B*, and *SOX17* in the TO cultures.^17^ Importantly, the bulk RNA-sequencing analysis recapitulated our initial findings that the IL-27 cytokine (*p28, EBI3*) and receptor (*IL27RA, WSX-1*) subunits are constitutively expressed by TOs (**Figure 2D**). Our bulk RNA-sequencing analysis also revealed distinct transcriptional signatures of TOs cultured in the absence/presence of IL-27 and IFNλ (**Supplemental Table 2, Supplemental Figure 3**). Given the IL-27-mediated changes in gene expression observed in the basal state for vehicle and isotype controls, we instead sought to compare the transcriptional profile of IL-27-stimulated TOs to IL-27-neutralized TOs. This analysis revealed 10 genes that were significantly upregulated in IL-27-stimulated TOs relative to IL-27-neutralized TOs (**Figure 2E**). Of these 10 genes, we identified 5 common antiviral genes that were also significantly upregulated in IFNλ-stimulated TOs relative to IFNλ-neutralized TOs (*IFIT1, DTX3L, PARP9, PARP14, PSME1*; **Supplemental Table 3**). Finally, gene set enrichment analysis (GSEA) revealed an upregulation of viral regulatory pathways in both IL-27- and IFNλ-stimulated TOs (**Figure 2F**), with similar antiviral genes driving the enriched gene sets (**Supplemental Tables 4 and 5**). Overall, these data suggest that IL-27 functions to effectively induce antiviral gene expression in human trophoblasts and contributes to maintaining the basal antiviral state of the placenta. This study is the first to describe transcriptional responses to IL-27 signaling in the human trophoblast organoid model.

### IL-27 is primarily produced within the decidua region of the murine placenta

Previous studies have demonstrated constitutive production of IL-27 subunit p28 in human placental tissues throughout gestation.^13^ To characterize IL-27 production and kinetics at the murine maternal-fetal interface, we first obtained lysates from whole murine placentas including maternal decidua at various gestational timepoints and measured IL-27 protein levels via ELISA. We observed that IL-27 is constitutively produced in the murine placenta throughout gestation (**Figure 3A**). To visualize cellular production of IL-27 by maternal and fetal cells in the murine placenta, we next utilized p28-GFP reporter mice.^23^ We mated p28-GFP^-/-^ dams and p28-GFP^+/-^ sires to generate a litter of fetuses that either lacked (p28-GFP^Null^) or expressed fetal-derived p28-GFP (p28-GFP^F^). In parallel, we mated p28-GFP^+/-^ dams and p28-GFP^-/-^ sires to generate litters of fetuses that retained maternal p28-GFP expression only (p28-GFP^M^) or expressed both maternal- and fetal-derived p28-GFP (p28-GFP^FM^) (**Figure 3B**). Placentas were collected at embryonic day 13.5 (E13.5) for immunofluorescence microscopy, and the maternal decidua (Dc) and fetal labyrinth (Lb) regions were defined according to endothelial cell (CD31) architecture (**Figure 3C**). We observed robust p28-GFP signal within the decidua of p28-GFP^F,^ p28-GFP^M^, and p28-GFP^FM^ mice (**Figure 3D**). Interestingly, we observed significantly more fetal-derived p28-GFP than maternal-derived p28-GFP within the decidua (**Figure 3D and 3E**). Although we hypothesized that p28 would be expressed within the fetal labyrinth, as this region contains murine trophoblast lineages analogous to the human trophoblasts shown to constitutively express IL-27, we observed minimal p28-GFP signal within the labyrinth region of p28-GFP^F,^ p28-GFP^M^, and p28-GFP^FM^ mice (**Figure 3E**, **Supplemental Figure 4**).^6,24^ Together these findings indicate that IL-27 is primarily produced at the murine maternal-fetal interface within the decidua, largely by cells of fetal origin.

**Figure 3:**
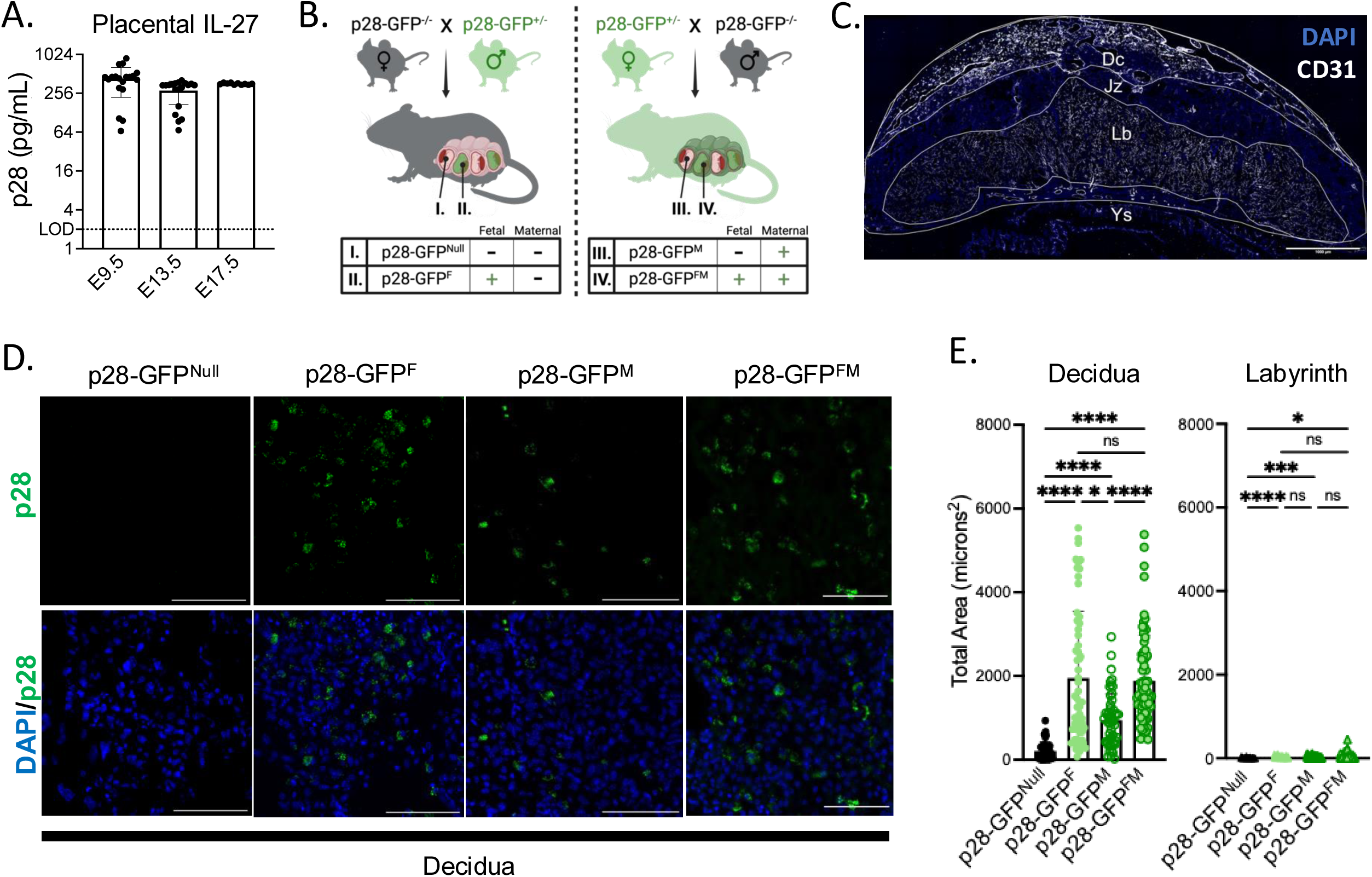
IL-27 is primarily produced within the decidua region of the murine placenta. **A.** Whole placental lysates (placenta and decidua) were collected from uninfected C57BL/6 mice at E9.5, E13.5, and E17.5 of gestation, and IL-27 protein levels were measured via ELISA. Graph represents placentas from 2 litters per timepoint. **B.** Schematic represents breeding schemes used to delineate maternal and fetal p28 production in the murine placenta and decidua. Left: *p28-GFP^-/-^* dam x *p28-GFP^+/-^* sire crosses were used to generate placental tissues that lacked (I., p28-GFP^Null^) or expressed fetal-derived p28-GFP (II., p28-GFP^F^). Right: *p28-GFP^+/-^* dam x *p28-GFP^-/-^* sire crosses were used to generate placental tissues that retained maternal p28-GFP expression only (III., p28-GFP^M^) or expressed both maternal- and fetal-derived p28-GFP (IV., p28-GFP^FM^). Graphics were created with Biorender.com. **C.** Representative immunofluorescent image of p28-GFP^F^ placenta at E13.5 of gestation. Yolk sac (Ys), labyrinth (Lb), junctional zone (Jz), and maternal decidua (Dc) regions were determined via immunofluorescence staining of endothelial cells (CD31; white) and nuclei (DAPI; blue). 20X magnification, stitched. Scale bar, 1000μm. **D.** Representative immunofluorescent images of p28-GFP within murine deciduas at E13.5. 10μm placental cross-sections were obtained for all conditions in Figure 3B and stained with anti-GFP (green) and DAPI (blue). Images representative of 4 placentas per condition. 20X magnification, zoomed. Scale bar 100μm. **E.** Quantification of total p28-GFP signal area (microns^2^) in decidua and labyrinth regions of placentas, as determined by ImageJ software. Graph represents measurements from 2-4 independent placentas per group, with 3-4 cross-sections obtained per placenta, and 4-8 images acquired per cross-section. Statistical analysis performed with Kruskal-Wallis ANOVA, *p<0.05, **p <0.001, ****p<0.0001.

### IL-27 signaling limits fetal pathology and placental ZIKV burdens during murine congenital infection

We observed that IL-27 regulates ZIKV infection in human trophoblasts *in vitro*, therefore we next sought to determine how placental IL-27 signaling influences pregnancy outcomes during broad inflammation and viral infection *in vivo*. To test this, we utilized immunocompetent human *STAT2* knock-in (hSTAT2 KI) mice, and hSTAT2 KI*IL27RA^-/-^ (IL27RA^-/-^) mice as a genetic disruption of IL-27 signaling.^25^ First, to trigger innate immune responses and induce a state of broad maternal inflammation in the absence of active viral replication, we challenged pregnant dams with high molecular weight polyinosinic-polycytidylic acid (HMW poly(I:C)), a synthetic double-stranded RNA and viral mimetic.^26^ We administered 20 mg/kg HMW poly(I:C) or PBS to pregnant hSTAT2 KI and IL27RA^-/-^ dams via intraperitoneal injection at E6.5. All dams were then euthanized at E13.5, a timepoint equivalent to the second trimester in humans, and fetal and placental tissues were collected for gross morphological analysis (**Figure 4A**). Importantly, we detected increased serum IL-6 levels in the HMW poly(I:C)-treated dams at 6 hours post-injection, indicating robust maternal immune activation as expected (**Figure 4B**). We then assessed fetal outcomes at the time of harvest, determining “healthy”, “early resorption”, or “total resorption” phenotypes for each fetus based on gross morphology and integrity of tissues, and quantified the proportion of fetal outcomes for each condition (**Figure 4C-D**). We observed that HWM poly(I:C)-stimulation of pregnant hSTAT2 KI mice resulted in some pathologic fetal outcomes (10.7% total resorption), however, this incidence was not significantly different in the absence of IL-27 signaling (3.6% early resorption, 10.7% total resorption), suggesting that IL-27 signaling does not protect against fetal pathology during broad maternal immune activation in pregnancy (**Figure 4C**).

**Figure 4:**
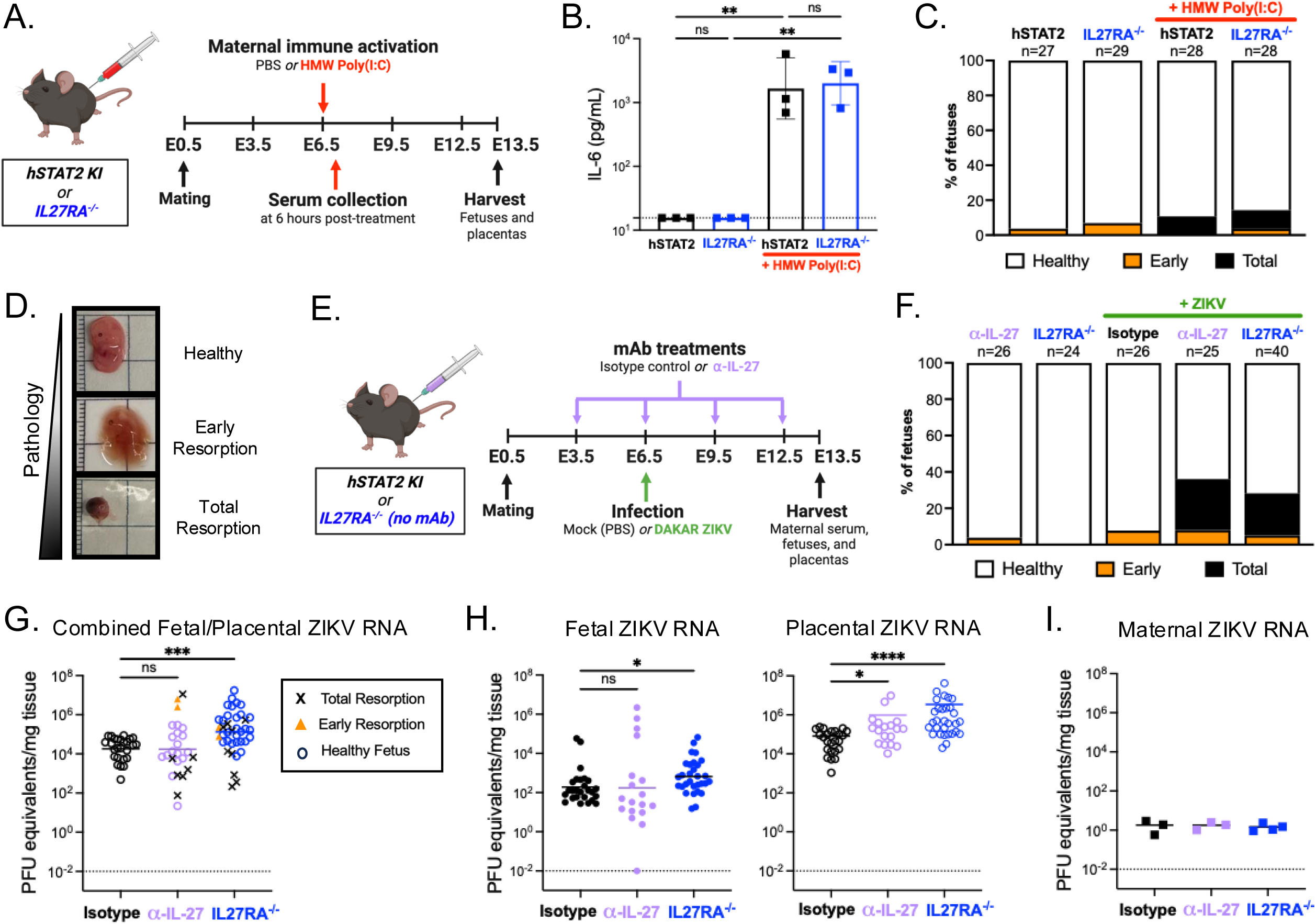
IL-27 signaling limits fet al pathology and pla cental ZIKV burdens during murine congenital inf ection. **A.** Schematic displaying the timeline for mating and treatment of human STAT2 knock-in (hSTAT2 KI) mice with high molecular weight polyinosinic-polycytidylic acid (HMW poly(I:C)). Pre gnant, hSTAT2 KI and IL27RA^-/-^ dams were treated with HMW poly(I:C) or PBS via intrape rito neal injection at E6.5. Maternal serum was collecte d via submandib ular (cheek) blee d at 6 hou rs post-injection. All dams were euthan ized at E13.5 and fetuses and placentas were collecte d for gro ss morph ology analysis. Gr aphics were created with Bior ender.com. **B.** Ser um IL-6 levels in PBS- and HMW Poly(I:C)-treated dams at 6 hou rs post-injection as determine d via ELISA. n=3 dams per gro up. Statistical ana lysis per for med via ord inary two-way ANOVA, **p<0.01. **C.** Pro portion of fetuse s exhibiting hea lthy, ear ly resorp tion , or total resorp tion phe notype s at E13.5 in PBS- and HMW Poly(I:C)-treated dams. Gr aph displays total numbe r of fetuse s from 3-4 litte rs per condition. **D.** Representative images dep icting fetal outcomes at E13.5. “Healthy”, “ea rly resorp tion ”, or “total resorp tion ” phe notype s were determine d for each fetus based on gro ss morph ology and integrity of tissues. Intact embryos lacking any visib le signs of patholo gy were characterized as “he althy.” Embryos with a distinguishable fetus and placenta but some loss in tissue integrity were characterized as “ea rly resorption.” Visibly smaller embryos that lacked a clearly defined fetus and placenta due to hemor rhage and tissue necrosis were characterized as “total resorp tion.” **E.** Schematic displaying the timeline for mating and infectio n of hSTAT2 KI mice. Pre gnant, hSTAT2 KI mice were treated with isotype control or α-IL-27 antibod y via intrape rito neal injection at E3.5, E6.5, E9.5, and E12.5, and infecte d with 6.75×10^5^ PFU mouse-ada pted DAKAR ZIKV or PBS at E6.5 via footpad injection. Add ition ally, pre gnant IL-27 receptor knocko ut (IL27RA^-/-^) mice were infecte d with 6.75×10^5^ PFU mouse-ada pted DAKAR ZIKV or PBS at E6.5 via footpad injection. All dams were euthan ized at E13.5 and fetuse s, placentas, and maternal serum were collecte d for further ana lysis. Gr aphics were created with Bior ender.com. **F.** Pro portion of fetuse s exhibiting hea lthy, ear ly resorption, or total resorp tion phe notype s at E13.5 in PBS- and ZIKV-infecte d dams. Gr aph displays total numbe r of fetuse s from 3-4 litte rs per condition. **G.** Fetal and placental ZIKV bur dens at E13.5 were determine d via qua ntitative RT-PCR. Each data poin t rep resents the combined ZIKV bur den of one matched fetus and placenta, nor ma lized to combined tissue weight. Sha pe of data poin t rep resents observed fetal phe notype. Statistical ana lysis per for med via Kru skal-Wallis ANOVA, ***p<0.001, ns= not significa nt. **H.** Left: Gr aph rep resents fetal ZIKV bur dens, nor ma lized to fetal weights. “Healthy” fetal phenotypes plotted. Right: Gr aph rep resents placental ZIKV bur dens, nor ma lized to placental weights. Placentas match ed to fetuse s with hea lthy phe notype s plotted. All gra phs rep resent 3-4 litte rs per condition. Statistical ana lysis per for med via Kru skal-Wallis ANOVA , *p<0.05, **p<0.01, ***p<0.001, ****p<0.000 1, ns= not significa nt. **I.** Circulating ZIKV RNA in maternal serum at E13.5 was measured via qua ntitative RT-PCR and nor ma lized to serum weight. n=3-4 dams per gro up. Not significa nt via Kru skal-Wallis ANOVA. **H.** Pro portion of fetuse s exhibiting hea lthy, ear ly resorp tion , or total resorp tion phe notype s at E13.5. Gr aph displays total numbe r of fetuse s from 3-4 litte rs per condition.

We next aimed to define how placental IL-27 signaling influences congenital viral infection. We obtained IL-27-deficient mice via antibody neutralization or genetic disruption of IL-27 signaling. For antibody neutralization, pregnant hSTAT2 KI dams were administered α-IL-27 or isotype control antibody via intraperitoneal injection beginning early in gestation and infected with mouse-adapted DAKAR ZIKV at E6.5 via footpad injection.^25^ As a genetic disruption of IL-27 signaling, pregnant IL27RA^-/-^ dams were also infected with mouse-adapted ZIKV at E6.5. Dams were euthanized at E13.5 and maternal serum, placentas, and fetuses were collected for further analysis (**Figure 4E**). In the absence of infection, the fetuses of IL-27-neutralized and IL27RA^-/-^ dams exhibited healthy outcomes and no observable pathology at E13.5 (**Figure 4F**). During congenital ZIKV infection, a small proportion of early resorption phenotypes were observed in the fetuses of isotype control-treated dams at E13.5 (7.7%; **Figure 4F**). However, there was a significantly greater incidence of resorption in the fetuses of IL-27-neutralized (28% total resorption; 8% early resorption) and IL27RA^-/-^ (23.1% total resorption; 5.1% early resorption) dams during congenital ZIKV infection (**Figure 4F, Supplemental Figure 5, Supplemental Table 6**).

We next measured ZIKV RNA levels in the fetuses, placentas, and maternal serum via quantitative RT-PCR. To include all fetuses in our analysis of viral burden, including those exhibiting the “total resorption” phenotype that prevents accurate separation of the placenta and fetus, we first evaluated the combined fetal and placental ZIKV burdens for each conceptus. We observed significantly higher ZIKV burdens in the combined fetal/placental tissues from IL27RA^-/-^ dams, suggesting that IL-27 may play a role in limiting ZIKV infection at the maternal-fetal interface (**Figure 4G**). When considering the intact fetuses alone, we found no significant difference in ZIKV burdens in the fetuses of control, IL-27-neutralized, and IL27RA^-/-^ dams, suggesting that fetal pathology is independent of ZIKV burden (**Figure 4H**). Interestingly, we observed significantly higher ZIKV burdens in the placentas of IL-27-neutralized and IL27RA^-/-^ dams relative to control placentas (**Figure 4H**). Finally, we found that the fetal pathology phenotypes were not driven by maternal ZIKV burdens, as the viral RNA was effectively cleared in the maternal serum as expected (**Figure 4I**). Collectively, these data indicate that IL-27 signaling is required to prevent fetal pathology and can restrict viral replication in placental tissues during congenital ZIKV infection.

## DISCUSSION

With this study, we sought to dive into the complex cytokine signaling mechanisms that facilitate immunological crosstalk between mother and fetus in the placenta and explore the previously uncharacterized role of placental IL-27 signaling. Using a primary human trophoblast organoid model, we found that IL-27 signaling machinery is expressed by fetal trophoblasts, and that IL-27 is capable of restricting ZIKV infection of trophoblasts similarly to IFNλ (**Figure 1**). Type III IFNs are a known family of antiviral cytokines produced by the placenta that have been shown to limit ZIKV infection of trophoblasts through antiviral gene expression mediated by autocrine signaling.^10–12^ Notably, our GSEA analysis of bulk RNA-sequencing data revealed similar transcriptional profiles of IL-27- and IFNλ-stimulated trophoblast organoids, with overlapping changes in antiviral gene expression, albeit to differing degrees (**Figure 2**). These findings underscore the numerous lines of defense that contribute to the robust antiviral state of the placenta during gestation.

IL-27 and IFNλ signaling interplay has been previously described in other contexts, with one study finding that IL-27 can drive IFNλ activation in virus-infected liver cells.^27^ To directly investigate cytokine signaling interplay in the placenta during congenital infection, future studies could explore ZIKV infection outcomes in the absence of both IL-27 and IFNλ. One current limitation of our studies comes from the ability to capture an unstimulated TO state, due to constitutive cytokine production in primary trophoblast organoids (**Figure 1**). Genetic manipulation of TO lines has been a technical challenge due to their derivation from primary human tissues; however, ongoing studies are pioneering methods for the genetic modification of TO lines which would allow for further dissection of the antiviral signaling interplay of these two cytokine families in trophoblasts *in vitro*.

In our murine model of congenital ZIKV infection, we observed that IL-27 signaling restricted placental ZIKV burdens and was protective against pathologic fetal outcomes indicating that IL-27 was critical to infection at gestational day E6.5 (**Figure 4**). Curiously, one study suggests that placental IFNλ signaling has both protective and pathogenic effects during murine congenital ZIKV infection, depending on the gestational timepoint.^28^ When pregnant dams were challenged with infection early in pregnancy (E7) IFNλ was pathogenic, whereas IFNλ was protective against infection later in pregnancy (E.9).^28^ Future studies are needed to explore the possibility of gestational-age dependent effects of IL-27 signaling during congenital murine ZIKV infection. Additionally, it is possible that IFNλ and IL-27 are functioning with varying mechanisms at the maternal-fetal interface during different timepoints of gestation. Future studies could elucidate the interplay of IL-27 and IFN-λ signaling at mid-gestation, and their distinct and shared contributions to protective antiviral immunity.

Though our study primarily explores the consequences of IL-27 signaling in placental trophoblasts, the placenta and decidua are both comprised of diverse cellular immune components, with many IL-27-responsive immune cells present in these tissues. For example, T cells, macrophages, natural killer cells, B cells and dendritic cells are all present at the maternal-fetal interface beginning early in gestation and play a key role in mediating a healthy pregnancy.^6,29^ IL-27 is known to potently regulate each of these subsets beyond the context of pregnancy, and prior studies show that IL-27 broadly restricts CD4+ T cells and dampens Th1, Th2, and Th17 cell-mediated pathology during acute respiratory viral infections.^30–33^ Yet, it remains unclear how IL-27 signaling contributes to immune cell regulation at the maternal-fetal interface. We observed increased tissue pathology in the fetuses of IL-27 deficient dams during congenital ZIKV infection, but not in the fetuses of IL-27-deficient dams treated with HMW poly(I:C) (**Figure 4**); however, these experiments did not explore the possibility of IL-27 regulating immune cell-mediated pathology. Thus, the role of IL-27 in regulating immune cell activity specifically during congenital ZIKV infection warrants further investigation.

Collectively, our work provides new insight into the functional role of IL-27 at the maternal-fetal interface and identifies IL-27 as a protective antiviral cytokine in the placenta. IL-27 has distinct protective roles in the context of other viral and microbial infections such as HIV-1, Influenza A virus (IAV), and *Toxoplasma gondii*, however this study is the first to demonstrate protective IL-27 activity at the maternal-fetal interface.^22,32,34–37^ Given our use of ZIKV as a model of congenital infection, future studies are needed to evaluate whether placental IL-27 signaling broadly contributes to protective immunity against other congenital pathogens such as Cytomegalovirus, *Toxoplasma gondii*, and *Listeria monocytogenes*. In addition to ZIKV, there are many other emerging and re-emerging viruses that pose a major threat to global health and to highly susceptible populations such as pregnant individuals and developing fetuses. We show that IL-27 signaling basally induces antiviral gene (*IFIT1, MX1, OAS2*) expression and viral regulation pathways in fetal trophoblasts (**Figure 2**), thus suggesting possible mechanisms through which IL-27 signaling could restrict other viruses known to infect fetal trophoblasts such as Rift Valley Fever virus, and recently emerging viruses with the capacitary for congenital infection, such as Oropouche virus.^38–42^ Finally, our study highlights therapeutic potential for IL-27 in the context of pregnancy and congenital infection. Recent studies have demonstrated the ability to specifically deliver mRNA to the placenta through ionizable lipid nanoparticle (LNP) technology to treat placental disorders such as pre-eclampsia.^43^ Future investigations could focus on applying the cutting-edge mRNA-LNP platform to effectively deliver IL-27 to the placenta and enhance antiviral responses *in vivo*.

## METHODS

### Human placental tissue samples

Placental villi and decidua were obtained from first trimester (6-12 post-conception weeks) pregnancy terminations performed at the University of Pennsylvania Family and Pregnancy Loss Center (IRB#827072). All subjects were counseled appropriately and provided written informed consent. Patients with preexisting medical conditions, pregnancy complications, or maternal infection were excluded from the study. Collected tissues were stored on ice and organoid derivation was carried out within 1 hour of procurement.

### Derivation and culture of trophoblast and decidual organoids

Primary trophoblast organoid (TO) and decidual organoid (DO) cultures were derived from first trimester placental tissues following methods previously established by Turco et al.^17–19^ Briefly, placental villi and decidual tissues were separated and washed in Ham’s/F12 media (Corning, MT10-080-CV) supplemented with 1% Penicillin/Streptomycin (ThermoFisher, 15140122) to remove blood. Villous tissues were scraped with a scalpel and enzymatically digested to isolate villous trophoblast stem/progenitor cells, while decidual tissues were minced with a scalpel and enzymatically digested to isolate endometrial gland stem/progenitor cells. TO progenitor cells were seeded in 25µl Matrigel domes, and plated with 250µl trophoblast organoid media (TOM) consisting of Advanced DMEM/F12 (ThermoFisher, 12634-010) supplemented with 1x B27 (ThermoFisher, 17504-044), 1x N2 (ThermoFisher, 17502-048), 2mM GlutaMAX (ThermoFisher, 35050-061), 100µg/mL Primocin (InvivoGen, ant-pm-1), 1.25mM N-acetyl-L-cysteine (Sigma-Aldrich, A9165), 5µM Y-27632 (Sigma-Aldrich, Y0503), 50ng/mL recombinant human EGF (Peprotech, AF-100-15), 100ng/mL recombinant human FGF2 (Peprotech, 100-18B), 80ng/mL recombinant human R-spondin 1 (R&D Systems, 4645-RS-01M/CF), 50ng/mL recombinant human HGF (Peprotech, 100-39), 1.5µM CHIR99021 (Tocris, 4423), 500nM A83-01 (Tocris, 2939), and 2.5µM Prostaglandin E2 (Tocris, 2296). DO progenitor cells were seeded in 25µl

Matrigel domes, and plated with 250µl expansion media (ExM) consisting of Advanced DMEM/F12 (ThermoFisher, 12634-010) supplemented with 1x B27 (ThermoFisher, 17504-044), 1x N2 (ThermoFisher, 17502-048), 2mM GlutaMAX (ThermoFisher, 35050-061), 100µg/mL Primocin (InvivoGen, ant-pm-1), 1.25mM N-acetyl-L-cysteine (Sigma-Aldrich, A9165), 50ng/mL recombinant human EGF (Peprotech, AF-100-15), 100ng/mL recombinant human FGF2 (Peprotech, 100-18B), 80ng/mL recombinant human R-spondin 1 (R&D Systems, 4645-RS-01M/CF), 50ng/mL recombinant human HGF (Peprotech, 100-39), 500nM A83-01 (Tocris, 2939), 10mM Nicotinamide (Sigma-Aldrich, N0636), and recombinant human NOGGIN (Peprotech, 120-10C). TO and DO cultures were routinely passaged every 6-8 days, with full media changes every 2-3 days, and were incubated at 37°C with 5% CO_2_. TOs and DOs used in this study were derived from 3 independent placentas (**Supplemental Table 1**). Sex of placentas and resulting TO lines were determined via quantitative RT-PCR using primers for *RPS4Y1* (Taqman, Hs00606158_m1; male placentas) and *Xist* (Taqman, Hs01079824_m1; female placentas).

### Enzyme Linked Immunosorbent Assays (ELISA)

Conditioned media (CM) was collected from TO and DO cultures at 1, 24, 48, and 72 hours post-passage. Secreted hCG was detected in organoid CM using One Step Pregnancy Test strips (Accumed, HCG25-PP). CM hCG levels were also measured via hCG ELISA (R&D Systems, DY9034) to validate TO and DO lines prior to experimentation. IL-27 ELISA (R&D Systems, DY2526) was performed according to manufacturer’s protocol using CM from 3-5 replicate TO wells for 3 independent TO donors, and a 2-fold serial dilution standard curve of recombinant human IL-27 plated in technical duplicate. For mouse IL-27 ELISAs, pregnant hSTAT2 KI mice were euthanized at gestational days E9.5, E13.5, and E17.5. Whole placental tissues, including maternal decidua, were collected and homogenized, and a mouse IL-27 p28 ELISA (R&D Systems, M2728) was then performed according to manufacturer’s protocol. Individual placental lysates and a 2-fold serial dilution standard curve of recombinant mouse IL-27 p28 were plated in technical duplicate. To validate maternal immune activation by HMW poly (I:C), an in-house IL-6 ELISA (Biolegend, 504501/504601) was performed on mouse serum samples. Blood was collected from HMW poly(I:C)- and control-treated dams via submandibular (cheek) bleed at 6 hours post-treatment, centrifuged to remove cells, and the remaining serum was collected (Sarstedt, 41.1378.005). IL-6 ELISAs were performed using serum from 3-4 dams per group, plated in technical duplicate. A standard curve of recombinant mouse IL-6 (Biolegend, 575702) was prepared according to manufacturer’s instructions and plated in duplicate. All TO and mouse ELISA absorbance readings were determined via plate reader at 450nm and converted to concentration using Microsoft Excel software.

### Immunofluorescence microscopy

Murine placentas and TOs were fixed in 4% paraformaldehyde overnight at 4°C prior to embedding in OCT (Sakura, 4583) and sectioning (8-10μm) with a Leica Cryostat. For all immunofluorescence experiments, slides were washed three times with 1x PBS prior to membrane disruption with PBS supplemented with 0.4% Triton X-100 (PBST; Sigma-Aldrich, T8787). All slides were blocked with 2% normal goat serum (Sigma-Aldrich, G9023-5ML) in PBST for one hour at room temperature prior to overnight incubation with primary antibodies diluted in blocking solution at 4°C. Mouse placenta sections were incubated in primary antibodies against GFP (1:200, Invitrogen, A-21311) and PECAM-1/CD31 (1:10, Developmental Studies Hybridoma Bank, 2H8s), while TO sections were incubated in primary antibodies against IL-27RA (1:100, Bioss Antibodies, bs-2711R), SDC1 (1:200, Abcam, ab34164), and Ki67 (1:50, Invitrogen, 50-5698-82). Slides were subsequently washed three times with PBST. After washing, slides were incubated with secondary antibodies (Jackson ImmunoResearch) and 1x DAPI (ThermoFisher, AC20271010) diluted in PBST for 2 hours at room temperature. Following incubation, mouse placenta slides were incubated in TrueBlack Lipofuscin Autofluorescence Quencher (Biotium, 23007) for three minutes at room temperature to reduce background signal intensity. All slides were washed with PBST and covered with mounting media (ThermoFisher, P36961) and a coverslip prior to imaging (Eclipse Ti2, Nikon). Immunofluorescence images were processed using NIS-Elements and ImageJ software. Total area of p28-GFP signal in murine placenta images was quantified using the ImageJ particle count application.

### Virus

For TO infections, ZIKV Cambodian FSS13025 strain (ZIKV-CAM) was obtained from the World Reference Center for Emerging Viruses and Arboviruses at University of Texas Medical Branch, Galveston. For *in vivo* infections, mouse-adapted ZIKV DAKAR was obtained from Michael Diamond’s laboratory at Washington University in St. Louis. ZIKV-CAM stocks were propagated in C6/36 mosquito cells (*Aedes albopictus*) (ATCC, CRL-1660), while ZIKV DAKAR stocks were grown in Vero cells (ATCC, CCL-81). C6/36 cells were maintained in Dulbecco’s Modified Eagle’s Medium (ThermoFisher, 11965092) supplemented with 10% heat-inactivated fetal bovine serum (ThermoFisher, A5670701), 1% penicillin/streptomycin (ThermoFisher, 15140122) and 1% tryptose phosphate broth (Sigma-Aldrich, T8159) at 30°C. Vero cells were maintained at 37°C in Dulbecco’s Modified Eagle’s Medium supplemented with 10% heat-inactivated fetal bovine serum and 1% penicillin/streptomycin. Cell lines were authenticated by morphology and were routinely tested for mycoplasma contamination. For high-titer doses, ZIKV was concentrated by ultrafiltration (Centricon Plus-70; MWCO: 30,000).

### Trophoblast organoid infection and stimulation

Prior to all TO experiments, TOs were routinely passaged and plated in Matrigel. For TO infections, TOs were treated with either isotype control (R&D, AB-108-C; R&D, MAB003), 10μg/mL α-IL-27 (R&D, AF2526) or 4μg/mL α-IFNλ1 (R&D, MAB15981) and 4μg/mL α-IFNλ3 (R&D, MAB5259) antibodies in fresh TOM at days 3 and 5 post-passage. At day 5 post-passage, TOs were removed from Matrigel domes and infected with 10^5^ PFU ZIKV-CAM or PBS in fresh TOM at 37°C and 5% CO_2_ for 2 hours. After 2 hours, virus was washed from TO cultures and replaced with fresh TOM and antibodies. At 24 hours post-infection, RNA was isolated from organoids and ZIKV burdens were determined via quantitative RT-PCR. Schematic of TO infection timeline included in **Supplemental Figure 2**. For **Figures 2B and 2C**, Isotype-treated TOs were administered PBS and isotype control antibody (R&D, AB-108-C; R&D, MAB003) in fresh TOM at days 3 and 5 post-passage. Rux-treated TOs were administered 20μM Ruxolitinib (Stem cell, 73404) and isotype control antibody in fresh TOM at days 3 and 5 post-passage. IFNλ and IL-27-treated TOs were treated with 10μg/mL α-IL-27 (R&D, AF2526) or 4μg/mL α-IFNλ1 (R&D, MAB15981) and 4μg/mL α-IFNλ3 (R&D, MAB5259) antibodies in fresh TOM at days 3 and 5 post-passage. At day 5 post-passage, IFNλ and IL-27-treated TO cultures were then stimulated with 100ng/mL each of recombinant human IFNλ1 (R&D, 1598-IL-025/CF) and recombinant human IFNλ3 (R&D, 5259-IL-025/CF), or 500ng/mL recombinant human IL-27 (Peprotech, 200-38) in fresh TOM, respectively. At 24 hours post-stimulation, all organoids were collected for RNA isolation and quantitative RT-PCR. For bulk RNA sequencing experiments, TOs were treated with either isotype control 10μg/mL α-IL-27 or 4μg/mL α-IFNλ1 and 4μg/mL α-IFNλ3 antibodies in fresh TOM at days 3 and 5 post-passage. At day 5 post-passage, additional TO cultures were stimulated with either vehicle control (PBS), 100ng/mL each of recombinant human IFNλ1 and IFNλ3, or 500ng/mL recombinant human IL-27 in fresh TOM. At day 6 post-passage, all organoids were collected for RNA isolation and sequencing.

### RNA extraction, cDNA generation, and quantitative RT-PCR

TOs and DOs were collected at 6-8 days post-passage for RNA analysis, depending on their size and density. To ensure RNA yield, 2-4 Matrigel domes containing organoids from a single donor were pooled in 400µl TRIzol reagent (Invitrogen, 15596026) for RNA analysis. Organoid RNA was purified via phenol-chloroform extraction and ethanol precipitation, using Phase Lock Gel Heavy tubes (QuantaBio, 2302830) according to manufacturer’s instructions. RNA quality and concentration were determined using a Thermo Fisher Scientific Nanodrop One spectrophotometer. Total RNA was reverse transcribed using iScript cDNA synthesis kit (Bio-Rad). Quantitative RT-PCR was performed using SYBR Green Master Mix reagent (Thermo Fisher Scientific, A25742) and human gene-specific primers (**Supplemental Table 4**). All samples were run in technical duplicate on a QuantStudio 3 Real-Time PCR System. Gene expression of *p28*, *EBI3*, *WSX1*, and *gp130* were determined based relative to housekeeping gene *HPRT1* expression (ΔCT method). For TO infections, ZIKV E expression was normalized to the isotype internal reference via the ΔΔCT method.

For RNA extraction and quantification of murine viral loads, tissues were collected in 600μl TRIzol reagent and homogenized (MP Biomedical, 116004500). Homogenates were then centrifuged for 5 minutes at 10,000RPM, and 100μl of the clarified homogenate was then diluted with 300μl TRIzol. An equal volume of 100% ethanol was then added to the diluted sample for RNA extraction. Viral RNA was extracted using a Direct-zol RNA Miniprep kit (Zymo Research, R2050) according to manufacturer’s instructions. Reverse transcription and quantitative RT-PCR were performed as described above using mouse gene-specific primers (**Supplemental Table 4**). ZIKV genome equivalents were determined via a 100-fold serially diluted standard curve of ZIKV RNA extracted from viral stock. Total ZIKV genome equivalents per tissue were then standardized to the tissue weight in mg.

### Bulk RNA-sequencing

RNA was isolated from TOs for bulk RNA-sequencing using the Direct-zol RNA Miniprep kit (Zymo Research, R2050) according to manufacturer’s instructions. RNA was then processed for bulk RNA-sequencing at the Children’s Hospital of Philadelphia High Throughput Sequencing Core. Sequencing libraries were prepared using Illumina’s stranded total RNA Prep kit with Ribo-Zero and the NovaSeq 6000 SP Reagent Kit v1.5 (200 cycles) (Illumina, #20040719). All data were analyzed using an adapted form of the open-source DIY transcriptomics lecture materials.^44^

### Mice

Wild-type (WT) C57BL/6J mice were purchased from The Jackson Laboratory (Strain #000664). Human STAT2 knock-in (hSTAT2 KI) mice (Gorman et. al. 2018) were obtained from Michael Diamond’s laboratory at Washington University in St. Louis. p28-GFP mice were obtained from Ross Kedl’s laboratory at the University of Colorado-Anschutz Medical Campus. All mice were maintained at the University of Pennsylvania under specific pathogen free conditions, and all experiments were performed in adherence to the University of Pennsylvania’s approved IACUC protocol. Mouse genotypes were determined using DNA extracted from maternal tail samples or fetal yolk sacs and previously described primers (**Supplemental Table 4**).^23,25^

### *In vivo* infections

Beginning at embryonic day 3.5 (E3.5), pregnant hSTAT2 KI dams were administered α-IL-27 (BioXCell, BE0326) or isotype control (BioXCell, BE0085) antibody via intraperitoneal injection, with subsequent treatments every three days (E6.5, E9.5, and E12.5). At E6.5, the control and IL-27-neutralized dams were then infected with 6.75×10^5^ PFUs mouse-adapted DAKAR ZIKV diluted in PBS via footpad injection. To mimic immune activation during viral infection, High Molecular Weight (HMW) Vaccigrade Poly(I:C) (Invivogen, vac-pic) was delivered to pregnant mice at E6.5 via intraperitoneal injection at 20mg/kg. All infected and HMW poly(I:C)-stimulated dams were euthanized at E13.5 and fetal, placental, and maternal tissues were collected for further analyses. All mouse infection and stimulation data are representative of 3-4 litters per treatment group.

#### Statistical analysis

Bulk RNA-sequencing data were analyzed in R studio using DIY transcriptomics lecture materials.^44^ All graphs were plotted and statistical analyses were performed using Microsoft Excel and GraphPad Prism software. Horizontal dashed lines are used on graphs to denote limits of detection (LOD) for the given assay. All data are expressed as mean +/-standard deviation (SD) or error of mean (SEM). Exact statistical test and number of biological samples (n) are detailed in the figure legends. A one-way ANOVA was used to determine significance when determining significance between multiple groups (>3). *p<0.05, **p<0.01, ***p<0.001, ****p<0.0001.

## DATA AVAILABILITY

The raw bulk RNA-sequencing data generated from TOs in this study have been submitted to the National Center for Biotechnology Information under BioProjectID #TBD. All original code will be deposited at Zenodo and will be publicly available as of the date of publication. Microscopy and sequencing data reported in this paper will be shared by the lead contact upon request.

## ACKNOWLEDGMENTS

We thank Dr. Michael Diamond from Washington University in St. Louis, MO, USA for providing the mouse adapted Zika DAKAR virus; Dr. Daniel Beiting at the University of Pennsylvania School of Veterinary Medicine, Philadelphia, PA, USA for assistance with bulk RNA sequencing analyses; Dr. Kevin R. Amses at the University of Pennsylvania, Philadelphia, PA, USA for assistance with biostatistical analyses; the High Throughput Sequencing (HTS) Core at the Children’s Hospital of Philadelphia (CHOP), PA, USA for assistance with bulk RNA sequencing experiments; Dr. Drake Philip at the University of Pennsylvania for assistance with trophoblast organoid analyses and passaging. This work was supported by the University of Pennsylvania Cell and Molecular Biology Training Grant T32 GM-07229 (MSM) and the National Institutes of Health IRACDA fellowship K12GM081259 (RLC). Additional funding provided by the Linda Pechenik Montague Investigator Award and the Pew Charitable Trusts Biomedical Scholars Program.

## AUTHOR CONTRIBUTIONS

M.S.M.: study conceptualization, study design, methodology, experimental work, data interpretation, and manuscript writing. B.B.: study design, experimental work and data interpretation. S.G.N.: methodology, RNA-sequencing data and bioinformatic analysis, manuscript review and editing. R.L.C.: methodology, manuscript review and editing. T.M-E.: methodology, manuscript review and editing. A.H.L.: methodology. S.M.: methodology. M.N.M.: study design, resources, and manuscript review. C.A.H.: study design, manuscript review and editing. K.A.J.: funding acquisition, study conceptualization, study design, methodology, manuscript writing, resources and supervision.

## REFERENCES

1. Megli, C. J. & Coyne, C. B. Infections at the maternal–fetal interface: an overview of pathogenesis and defence. Nat. Rev. Microbiol. 1–16 (2021) doi:10.1038/s41579-021-00610-y.

2. Coyne, C. B. & Lazear, H. M. Zika virus — reigniting the TORCH. Nat. Rev. Microbiol. 14, 707–715 (2016).

3. França, G. V. A. et al. Congenital Zika virus syndrome in Brazil: a case series of the first 1501 livebirths with complete investigation. The Lancet 388, 891–897 (2016).

4. Bhatnagar, J. et al. Zika Virus RNA Replication and Persistence in Brain and Placental Tissue. Emerg. Infect. Dis. 23, 405–414 (2017).

5. Hoen, B. et al. Pregnancy Outcomes after ZIKV Infection in French Territories in the Americas. N. Engl. J. Med. 378, 985–994 (2018).

6. Ander, S. E., Diamond, M. S. & Coyne, C. B. Immune responses at the maternal-fetal interface. Sci. Immunol. 4, eaat6114 (2019).

7. Sadler, A. J. & Williams, B. R. G. Interferon-inducible antiviral effectors. Nat. Rev. Immunol. 8, 559–568 (2008).

8. Yockey, L. J. et al. Type I interferons instigate fetal demise after Zika virus infection. Sci. Immunol. (2018).

9. Casazza, R. L., Lazear, H. M. & Miner, J. J. Protective and Pathogenic Effects of Interferon Signaling During Pregnancy. Viral Immunol. 33, 3–11 (2020).

10. Bayer, A. et al. Type III Interferons Produced by Human Placental Trophoblasts Confer Protection against Zika Virus Infection. Cell Host Microbe 19, 705–712 (2016).

11. Corry, J., Arora, N., Good, C. A., Sadovsky, Y. & Coyne, C. B. Organotypic models of type III interferon-mediated protection from Zika virus infections at the maternal–fetal interface. Proc. Natl. Acad. Sci. 114, 9433–9438 (2017).

12. Jagger, B. W. et al. Gestational Stage and IFN-λ Signaling Regulate ZIKV Infection In Utero. Cell Host Microbe 22, 366–376.e3 (2017).

13. Coulomb-L’Herminé, A. et al. Expression of interleukin-27 by human trophoblast cells. Placenta 28, 1133–1140 (2007).

14. Hunter, C. A. & Kastelein, R. Interleukin-27: Balancing Protective and Pathological Immunity. Immunity 37, 960–969 (2012).

15. Yoshida, H. & Hunter, C. A. The immunobiology of interleukin-27. Annu. Rev. Immunol. 33, 417–443 (2015).

16. Kwock, J. T. et al. IL-27 signaling activates skin cells to induce innate antiviral proteins and protects against Zika virus infection. Sci. Adv. 6, eaay3245 (2020).

17. Turco, M. Y. et al. Long-term, hormone-responsive organoid cultures of human endometrium in a chemically-defined medium. Nat. Cell Biol. 19, 568–577 (2017).

18. Turco, M. Y. et al. Trophoblast organoids as a model for maternal–fetal interactions during human placentation. Nature 564, 263–267 (2018).

19. Sheridan, M. A. et al. Establishment and differentiation of long-term trophoblast organoid cultures from the human placenta. Nat. Protoc. 15, 3441–3463 (2020).

20. Co, J. Y. et al. Controlling Epithelial Polarity: A Human Enteroid Model for Host-Pathogen Interactions. Cell Rep. 26, 2509–2520.e4 (2019).

21. Yang, L. et al. Innate immune signaling in trophoblast and decidua organoids defines differential antiviral defenses at the maternal-fetal interface. eLife 11, e79794 (2022).

22. Valdés-López, J. F. et al. Interleukin 27, like interferons, activates JAK-STAT signaling and promotes pro-inflammatory and antiviral states that interfere with dengue and chikungunya viruses replication in human macrophages. Front. Immunol. 15, (2024).

23. Kilgore, A. M. et al. IL-27p28 Production by XCR1+ Dendritic Cells and Monocytes Effectively Predicts Adjuvant-Elicited CD8+ T Cell Responses. ImmunoHorizons 2, 1–11 (2018).

24. Soncin, F., Natale, D. & Parast, M. M. Signaling pathways in mouse and human trophoblast differentiation: a comparative review. Cell. Mol. Life Sci. CMLS 72, 1291–1302 (2014).

25. Gorman, M. J. et al. An Immunocompetent Mouse Model of Zika Virus Infection. Cell Host Microbe 23, 672–685.e6 (2018).

26. Hameete, B. C. et al. The poly(I:C)-induced maternal immune activation model; a systematic review and meta-analysis of cytokine levels in the offspring. Brain Behav. Immun. -Health 11, 100192 (2020).

27. Cao, Y. et al. IL-27, a Cytokine, and IFN-λ1, a Type III IFN, Are Coordinated To Regulate Virus Replication through Type I IFN. J. Immunol. 192, 691–703 (2014).

28. Casazza, R. L., Philip, D. T. & Lazear, H. M. Interferon Lambda Signals in Maternal Tissues to Exert Protective and Pathogenic Effects in a Gestational Stage-Dependent Manner. mBio 13, e03857–21 (2022).

29. Vento-Tormo, R. et al. Single-cell reconstruction of the early maternal–fetal interface in humans. Nature 563, 347–353 (2018).

30. Villarino, A. V. et al. Positive and Negative Regulation of the IL-27 Receptor during Lymphoid Cell Activation. J. Immunol. 174, 7684–7691 (2005).

31. Findlay, E. G. et al. Essential Role for IL-27 Receptor Signaling in Prevention of Th1-Mediated Immunopathology during Malaria Infection. J. Immunol. 185, 2482–2492 (2010).

32. Liu, F. D. M. et al. Timed Action of IL-27 Protects from Immunopathology while Preserving Defense in Influenza. PLOS Pathog. 10, e1004110 (2014).

33. Muallem, G. et al. IL-27 Limits Type 2 Immunopathology Following Parainfluenza Virus Infection. PLOS Pathog. 13, e1006173 (2017).

34. Dai, L. et al. IL-27 inhibits HIV-1 infection in human macrophages by down-regulating host factor SPTBN1 during monocyte to macrophage differentiation. J. Exp. Med. 210, 517–534 (2013).

35. Swaminathan, S., Dai, L., Lane, H. C. & Imamichi, T. Evaluating the potential of IL-27 as a novel therapeutic agent in HIV-1 infection. Cytokine Growth Factor Rev. 24, 571–577 (2013).

36. Villarino, A. et al. The IL-27R (WSX-1) Is Required to Suppress T Cell Hyperactivity during Infection. Immunity 19, 645–655 (2003).

37. Aldridge, D. L. et al. Endogenous IL-27 during toxoplasmosis limits early monocyte responses and their inflammatory activation by pathological T cells. mBio 15, e00083–24.

38. McMillen, C. M. et al. Rift Valley fever virus induces fetal demise in Sprague-Dawley rats through direct placental infection. Sci. Adv. 4, eaau9812 (2018).

39. Oymans, J., Wichgers Schreur, P. J., van Keulen, L., Kant, J. & Kortekaas, J. Rift Valley fever virus targets the maternal-foetal interface in ovine and human placentas. PLoS Negl. Trop. Dis. 14, e0007898 (2020).

40. McMillen, C. M. & Hartman, A. L. Rift Valley Fever: a Threat to Pregnant Women Hiding in Plain Sight? J. Virol. 95, 10.1128/jvi.01394-19 (2021).

41. Megli, C. et al. Oropouche virus infects human placenta explants and trophoblast organoids. 2024.11.16.623866 Preprint at 10.1101/2024.11.16.623866 (2024).

42. Schwartz, D. A., Dashraath, P. & Baud, D. Oropouche Virus (OROV) in Pregnancy: An Emerging Cause of Placental and Fetal Infection Associated with Stillbirth and Microcephaly following Vertical Transmission. Viruses 16, 1435 (2024).

43. Swingle, K. L. et al. Placenta-tropic VEGF mRNA lipid nanoparticles ameliorate murine pre-eclampsia. Nature 637, 412–421 (2025).

44. Berry, A. S. F. et al. An Open-Source Toolkit To Expand Bioinformatics Training in Infectious Diseases. mBio 12, e0121421 (2021).

