## Supplemental Figures for "Interleukin-27 is antiviral at the maternal-fetal interface"

**Supplemental Figure 1:** Validation of first trimester TO and DO cultures

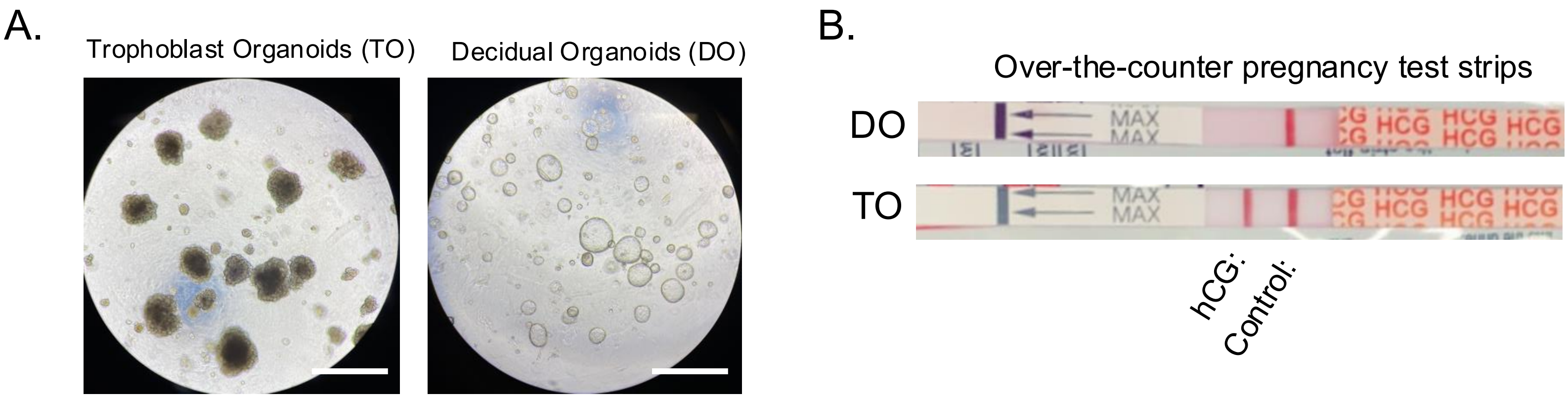


**Supplemental Figure 1. Validation of first trimester TO and DO cultures**

**A.** Representative brightfield images of TOs and DOs in culture. 5X magnification. Scale bar, 100µm.

**B.** Over the counter pregnancy test strips were used to detect secreted trophoblast-specific hormone hCG in TO- and DO-conditioned media.

**Supplemental Figure 2:** ZIKV infection of TO cultures

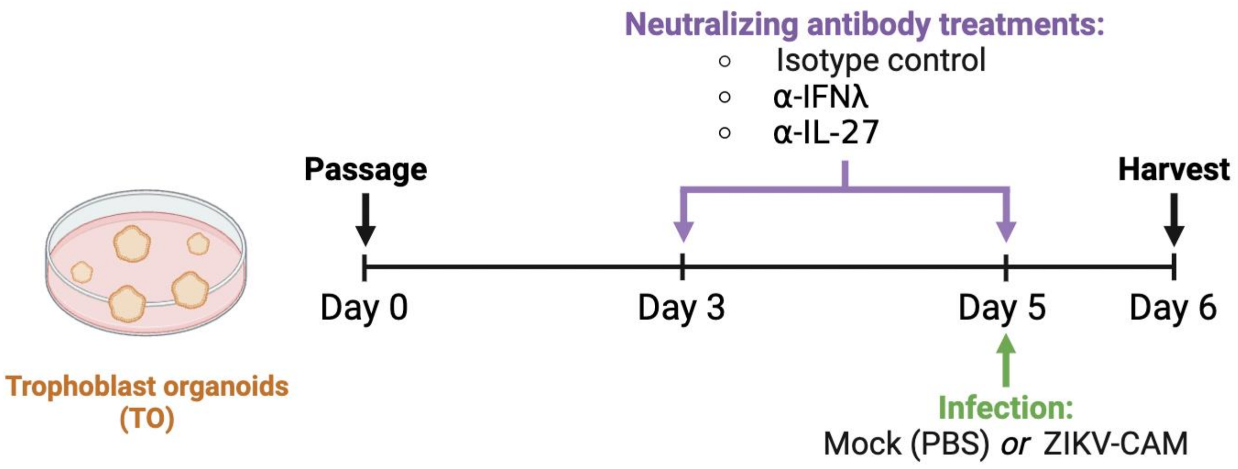


**Supplemental Figure 2. ZIKV infection of TO cultures**

**A.** Schematic of experimental design for TO infection with Cambodian ZIKV (ZIKV-CAMB) as quantified in **Figure 1E**. Graphics were created with Biorender.com.

**Supplemental Figure 3.** Bulk RNA-sequencing of TOs

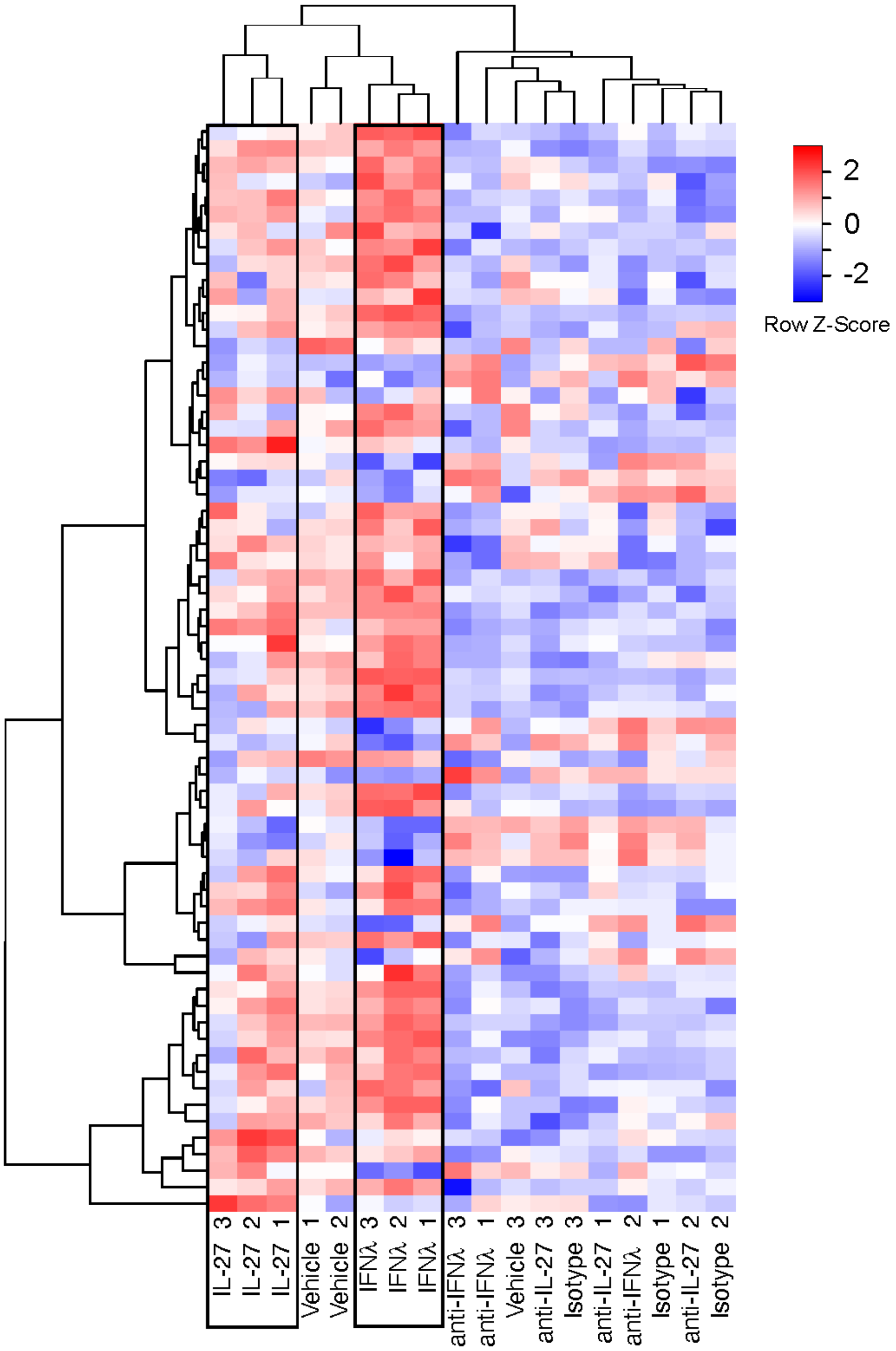


**Supplemental Figure 3. Bulk RNA-sequencing of TOs**

**A.** Heat map depicts gene expression data scaled by z-score for each row from bulk RNA sequencing of TOs. Columns represent samples clustered using Spearman correlation, and rows represent differentially expressed genes that were identified by comparing IL-27-stimulated to IL-27-neutralized TOs and IFNλ-stimulated to IFNλ-neutralized TOs, then clustered using Pearson correlation. 66 genes in the heatmap met cutoffs of p-value=0.05 and log fold-change=±0.5 and are listed in **Supplemental Table 2**. Data are representative of 3 independent samples per condition.

**Supplemental Figure 4.** p28 production within the murine labyrinth

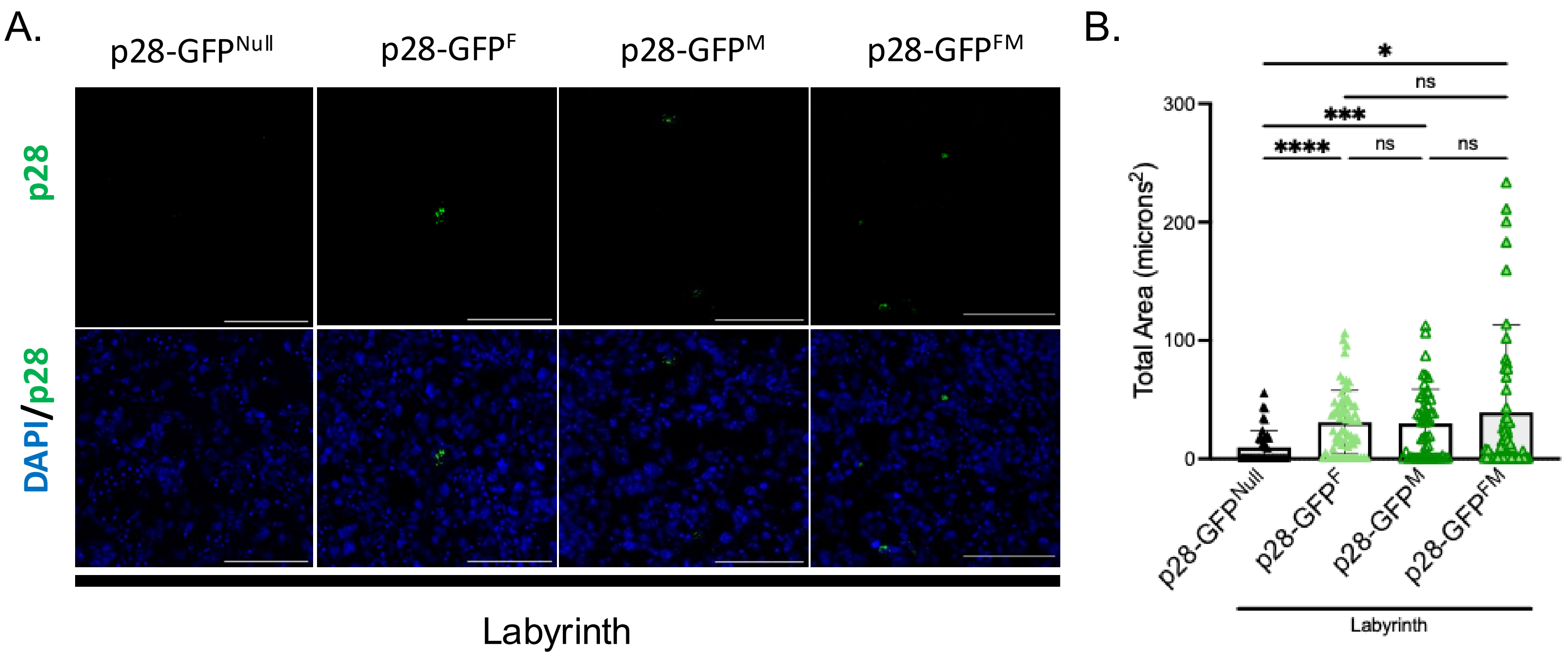


**Supplemental Figure 4. p28 production within the murine labyrinth**

**A.** p28-GFP localization within murine placental labyrinth at E13.5. 10µm placental cross-sections were stained with anti-GFP (green) and DAPI (blue) and imaged via microscopy. Representative images from 4 independent placentas. 20X magnification. Scale bar, 100µm. **B.** Quantification of total labyrinth p28-GFP signal area (microns$^2$), as shown in **Figure 3E**, adjusted for scale. Graph represents measurements from 2-4 independent placentas per group, with 3-4 cross-sections obtained per placenta, and 4-8 images acquired per cross-section. Statistical analysis performed with Kruskal-Wallis ANOVA, *p<0.05, **p <0.001, ****p<0.0001.

**Supplemental Figure 5.** IL-27 signaling limits fetal pathology during murine congenital ZIKV infection

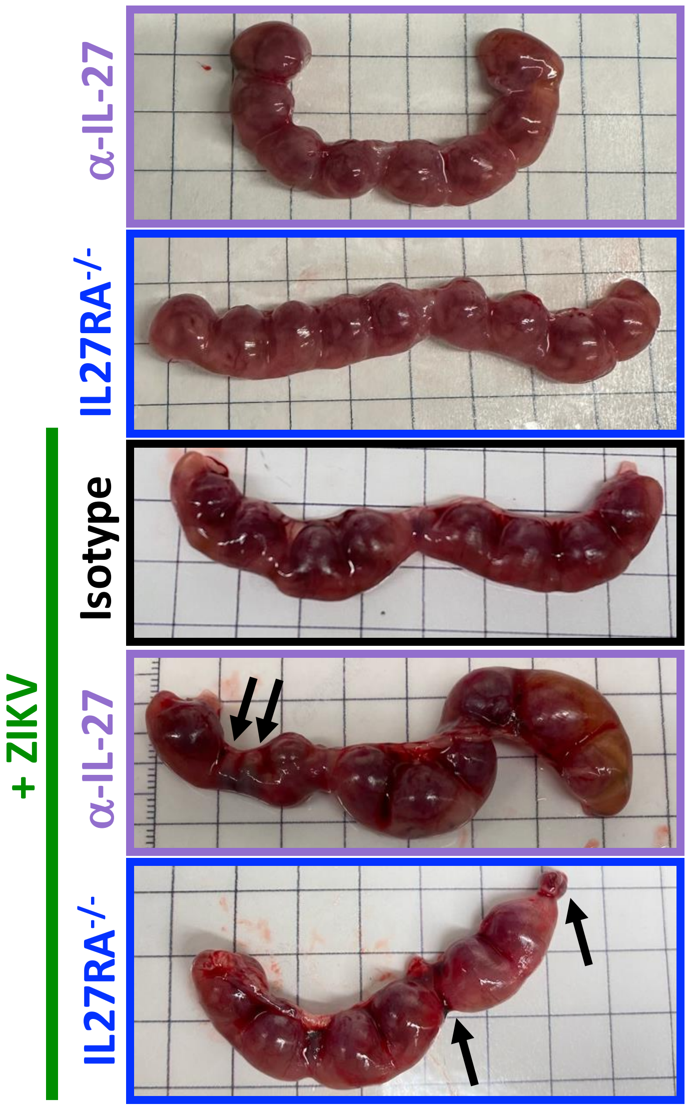


**Supplemental Figure 5. IL-27 signaling limits fetal pathology during murine congenital ZIKV infection**

**A.** Representative images of murine uterine horns isolated at E13.5 for tissue pathology and viral burden analyses. Black arrows highlight embryos exhibiting pathologic fetal outcomes.
